# The Development of Resistance to an Inhibitor of a Cellular Protein Reveals a critical interaction between the enterovirus protein 2C and a small GTPase Arf1

**DOI:** 10.1101/2023.05.09.539905

**Authors:** Ekaterina G. Viktorova, Samuel Gabaglio, Seyedehmahsa Moghimi, Anna Zimina, Bridge G. Wynn, Elizabeth Sztul, George A. Belov

## Abstract

The cellular protein GBF1, an activator of Arf GTPases (ArfGEF: Arf guanine nucleotide exchange factor), is recruited to the replication organelles of enteroviruses through interaction with the viral protein 3A, and its ArfGEF activity is required for viral replication. Here, we investigated the development of resistance of poliovirus, a prototype enterovirus, to increasing concentrations of brefeldin A (BFA), an inhibitor of GBF1. High level of resistance required a gradual accumulation of multiple mutations in the viral protein 2C. The 2C mutations conferred BFA resistance even in the context of a 3A mutant previously shown to be defective in the recruitment of GBF1 to replication organelles, and in cells depleted of GBF1, suggesting a GBF1-independent replication mechanism. Still, activated Arfs accumulated on the replication organelles of this mutant even in the presence of BFA, its replication was inhibited by a pan-ArfGEF inhibitor LM11, and the BFA-resistant phenotype was compromised in Arf1-knockout cells. Importantly, the mutations strongly increased the interaction of 2C with the activated form of Arf1. Analysis of other enteroviruses revealed a particularly strong interaction of 2C of human rhinovirus 1A with activated Arf1. Accordingly, the replication of this virus was significantly less sensitive to BFA than that of poliovirus. Thus, our data demonstrate that enterovirus 2Cs may behave like Arf1 effector proteins and that GBF1 but not Arf activation can be dispensable for enterovirus replication. These findings have important implications for the development of host-targeted anti-viral therapeutics.

## Introduction

Enteroviruses are small (+)RNA viruses of the genus *Enterovirus* of the family *Picornaviridae*. They comprise numerous established and emerging human pathogens such as Coxsackieviruses, poliovirus, rhinoviruses, enteroviruses D68 and A71, and many others. Enterovirus infections can be associated with diverse pathologies ranging from a routine common cold to life-threatening conditions including diabetes, myocarditis and fatal encephalitis (1). Yet, the current arsenal of anti-enteroviral tools is essentially limited only to anti-poliovirus and anti-enterovirus A71 vaccines (2, 3). The continuous research on anti-enterovirus therapeutics has so far produced limited results, and there are still no drugs officially approved to treat enteroviral infections (4). The major problem in the development of anti-enteroviral therapeutics is that these viruses rapidly develop resistance to any compounds targeting either viral proteins or cellular factors and pathways hijacked for viral replication (5). The latter ability runs against the expectations that overcoming the requirement for an essential cellular activity should represent a significant barrier for the virus, underscoring the limits of the current knowledge of the mechanistic details of viral replication.

In spite of antigenic diversity, enteroviruses share significant similarities in molecular mechanisms of replication, including the dependence on essential cellular factors. Poliovirus, a representative enterovirus and one of the best-studied animal viruses, serves as an important model for the investigation of enterovirus replication and virus-cell interaction. Poliovirus genome RNA of ∼7500nt serves both as an mRNA and as a template for replication. It codes for one polyprotein, which is processed co- and post-translationally by virus-encoded proteases into three structural and ten replication proteins, including stable intermediates of the polyprotein processing. Three of the non-structural viral proteins, 2B, 2C, and 3A have membrane-targeting domains that anchor the viral replication complexes on specialized membranous structures, replication organelles (6-8).

Cellular factors recruited to the replication organelles facilitate the development of their unique lipid and protein composition necessary for the optimal functioning of the viral replication machinery. The complete composition of the cellular proteins on the replication organelles, let alone their roles in the replication process, are far from understood. One of the cellular proteins important for the replication of enteroviruses as well as many others but not all (+)RNA viruses is GBF1, a guanine nucleotide exchange factor for small cellular GTPases of the Arf family (ArfGEF) (9-13). Arfs in their activated, GTP-bound form, associate with membranes and interact with multiple Arf effector proteins recruiting them to the membranes and regulating their activity. Hydrolysis of GTP results in the dissociation of the inactivated Arf-GDP from membranes. Human cells express five Arf isoforms (Arfs 1, 3, 4, 5, and 6), and the tight spatial and temporal control of the Arf activation-deactivation cycle on cellular membranes is important for the maintenance of the biochemical identity of cellular organelles and regulation of intracellular membrane trafficking (14, 15).

GBF1 is a large (∼200 KDa) multidomain protein that maintains a pool of activated Arfs at the ER-Golgi interface necessary for the functioning of the cellular secretory pathway. GBF1-dependent Arf activation is also important for membrane trafficking at the trans-Golgi network and delivery of certain proteins to lipid droplets (16-18). The ArfGEF activity of GBF1 is mediated by a centrally located Sec7 domain, while other domains of GBF1 contribute to the interaction of the protein with other cellular factors, its recruitment to membranes, and regulation of its activity (19-21). The Sec7 domains demonstrate a remarkable structural and functional conservation even among organisms of different kingdoms, yet the variances in amino-acid sequences make them differently susceptible to inhibitors (21-23). Among the 15 Sec7-containing proteins expressed in mammalian cells, ArfGEF activity of only GBF1, BIG1, and BIG2 can be inhibited by the fungal metabolite brefeldin A (BFA), while Golgicide A (GCA) inhibits GBF1 but not other ArfGEFs (24-31), and a recently developed compound LM11 inhibits the activity of all human ArfGEFs (32).

In cells infected with poliovirus and other enteroviruses, GBF1 interacts with the viral protein 3A and relocalizes to the replication organelles, which results in a massive accumulation of activated Arfs on replication membranes (12, 13, 33, 34). BFA and GCA strongly inhibit enterovirus replication, and such inhibition can be relieved only by the overexpression of GBF1 but not other ArfGEFs (12, 13, 35). Moreover, enteroviruses can efficiently replicate in cells expressing extensively truncated GBF1 mutants, but not those with non-functional Sec7 domains, indicating the importance of GBF1-driven Arf activation for replication (36). While the contribution of individual Arf isoforms to enterovirus replication remains poorly understood, it appears that activation of Arf1 plays the most important role in the development and/or functioning of the replication organelles (33).

Contrary to the expectations that inhibitors of cellular proteins such as GBF1 may pose a higher barrier to viral resistance, poliovirus mutants capable of replicating in the presence of BFA could be easily selected, at least at relatively low concentrations of the drug (37, 38). The resistance is conferred by single amino-acid substitutions in the membrane-targeted viral proteins 2C and 3A. The 3A mutations provided a major contribution to the resistance phenotype, and the combined effect of 2C and 3A mutations was required for the full level of resistance (37, 38). In the absence of BFA, replication of such a mutant is accompanied by a massive Arf activation, similar to the wt virus. Yet, in the presence of the drug, Arf activation could not be detected in a biochemical pull-down assay, which could be explained by either a lower level of replication, or a possible diminished requirement of the mutant for the activated Arf (39). Intriguingly, the BFA-resistant virus was still dependent on GBF1 both in the presence and in the absence of BFA, as evidenced by the sensitivity of replication to the siRNA-mediated knockdown of GBF1 expression (39). Moreover, the potency of BFA-resistant mutations was severely compromised in the context of a 3A mutation reducing the 3A-GBF1 interaction, suggesting that at least a residual amount of GBF1-activated Arf(s) is still required for the replication of the BFA-resistant poliovirus mutants (38).

In this study, we investigated the development of resistance of poliovirus replication to much higher concentrations of BFA than those used previously, with the goal of obtaining the most conspicuous phenotype that may help elucidate the mechanism of resistance. Importantly, such a selection was successful only after incremental increases of BFA concentrations. Sequencing the viral isolates from successive passages in increasing concentrations of BFA revealed dynamic changes in the mutational landscape of 2C and 3A. We further investigated the replication of the mutant that demonstrated the strongest BFA-resistant phenotype. Contrary to the previously observed pattern (37, 38), mutations in 2C but not in 3A were responsible for resistance to high level of BFA. However, the combination of mutations in 2C and 3A was required for BFA-dependent phenotype observed in some but not all cell types. The mutated 2C could confer a high level of BFA resistance even in the context of a 3A mutation previously shown to significantly inhibit 3A-GBF1 interaction (38), and in cells with knockdown of GBF1 expression. At the same time, replication of the resistant virus was accompanied by a robust accumulation of all Arf isoforms on the replication organelles in the absence and in the presence of BFA. We found that the replication of wt and BFA-resistant polio replicons is sensitive to the depletion of several BFA-insensitive ArfGEFs, suggesting that GBF1-independent Arf activation may contribute to the development and/or functioning of the replication organelles. Importantly, the BFA-resistant mutant was still sensitive to a pan-ArfGEF inhibitor LM11, and the BFA resistance was severely compromised by Arf1 knockout and to a lesser extent by Arf6 knockdown, indicating the need for Arf1 activation to sustain BFA-independent viral replication. Moreover, pull-down experiments revealed that the mutations in 2C conferring BFA resistance strongly increased 2C interaction with activated Arf1. Furthermore, our finding that 2C of rhinovirus 1A strongly interacts with Arf1 and that its replication is much less sensitive to BFA than that of poliovirus further supports the key role of the 2C interaction with activated Arf1 in BFA resistance. Together, our data strongly suggest that enterovirus 2C functions as an Arf effector protein and infer the mechanism of BFA resistance depending on the scavenging of activated Arf1 generated by cellular BFA-insensitive ArfGEFs by mutated 2C. Moreover, these results demonstrate that while the resistance of enterovirus replication to the inhibition of ArfGEF activity of GBF1 develops relatively easily, the requirement for activated Arfs could not be circumvented. Thus, the 2C-Arf interface could be a promising target for anti-viral therapeutics which would not only be refractable to the development of resistance, but also would be active only in infected cells as opposed to inhibitors such as BFA that target general cellular pathways.

## Materials and Methods

### Cells

Human cervical carcinoma HeLa cell line was a gift from Dr. Ellie Ehrenfeld (NIH). Human embryonic kidney HEK293 cell line and human colon cancer Caco2 cell line were a gift from Dr. Xiaoping Zhu (University of Maryland). Human hepatoma Huh7 cells were a gift from Dr. Liqing Yu (University of Maryland School of Medicine). Human osteosarcoma MG63 and human lung carcinoma A549 cells were purchased from ATCC. Stable HeLa cell lines expressing individual human Arfs fused to Venus fluorescent protein were described in (33). HeLa cells with CRISPR-CAS9-generated knockouts of Arf1, 3, 4, and 5 were described in (40) and were kindly provided by Dr. Martin Spiess (University of Basel, Switzerland). HeLa cells were grown in DMEM high glucose modification supplemented with pyruvate. HEK293, and Huh7 cells were grown in DMEM. MG63 cells were grown in Eagle’s MEM, and A549 cells were grown in F12K medium. All media were supplemented with 10% FBS.

### Selection of BFA-resistant polioviruses

Poliovirus type I Mahoney was propagated in HeLa cells, and the titer was determined by plaque assays. For the selection of resistant mutants, the first passage was performed by infecting a HeLa cell monolayer grown on a 35 mm dish at a multiplicity of infection (MOI) of 50 plaque-forming units (PFU) per cell. The virus adsorption was performed for 30 min at room temperature on a rocking platform in 0.5 ml of medium without the inhibitor. After adsorption, the cells were incubated in 2 ml of the complete growth medium with the indicated amount of BFA for 6 h at 37C. At 6 h p.i. the plate with cells and the medium was frozen. The virus was released by three cycles of freeze-thawing and the medium was clarified from cell debris by low-speed centrifugation. Half of the medium was used for the subsequent passaging, and the rest was used for plaque assays, viral RNA isolation, and sequencing.

### Plasmids

Plasmid pXpA-SH coding for the full-length poliovirus type I (strain Mahoney) cDNA with introduced SalI and HpaI restriction sites in the 5’ and 3’ non-translated regions was described in (41), and its derivative plasmid pXpA-RenR coding for a poliovirus replicon with Renilla luciferase gene substituting the capsid coding region was described in (42). To introduce the BFA-resistant mutations in 2C and 3A either PCR fragments synthesized using isolated RNA of BFA-resistant viruses as a template, or gene fragments synthesized by GeneArt service (Thermo Fisher) were introduced into pXpA-SH and pXpA-RenR using standard cloning techniques. All mutations were verified by sequencing. Plasmids p53CB3T7 and pACYC-RV1A coding for the cDNA of Coxsackievirus B3 (strain Nancy) and rhinovirus 1A (strain ATCC VR-1559) under the control of T7 promotor were kindly provided by Professor Frank van Kuppeveld (University of Utrecht, the Netherlands) and Dr. Margaret Scull (University of Maryland), respectively. Expression plasmids pM1-2Cs coding for 2Cs of poliovirus (wt and BFA-resistant), were constructed using pM1-MT vector for a high level of mammalian expression (Roche Biosciences). Cloning details are available upon request. Plasmids coding for human Arf1, 3, 4, 5, and 6 fused to EGFP were a gift from Dr. Catherine Jackson (Université Paris Diderot, France).

### Antibodies and reagents

Mouse monoclonal antibodies: anti-GBF1 was from BD Biosciences (Cat# 612116); anti-poliovirus 2C described in (43) was a gift from Professor Kurt Bienz, University of Basel, Switzerland, this antibody recognizes 2C of diverse enteroviruses; anti-β- actin antibody fused to horse radish peroxidase was from Sigma Aldrich (Cat# A3854). Rabbit polyclonal antibodies against GFP were from Abcam (Cat# ab290). GFP selector beads with a conjugated camelid single domain antibody against GFP were from NanoTag Biotechnologies (Cat. # N0310). These beads recognize multiple GFP derivatives including Venus. Secondary antibody conjugates with Alexa fluorofores were from Molecular Probes, those conjugated with horse radish peroxidase were from SeraCare. BFA was from Sigma Aldrich, LM11 was from Hit2Lead ChemBridge Chemical Store.

### siRNAs

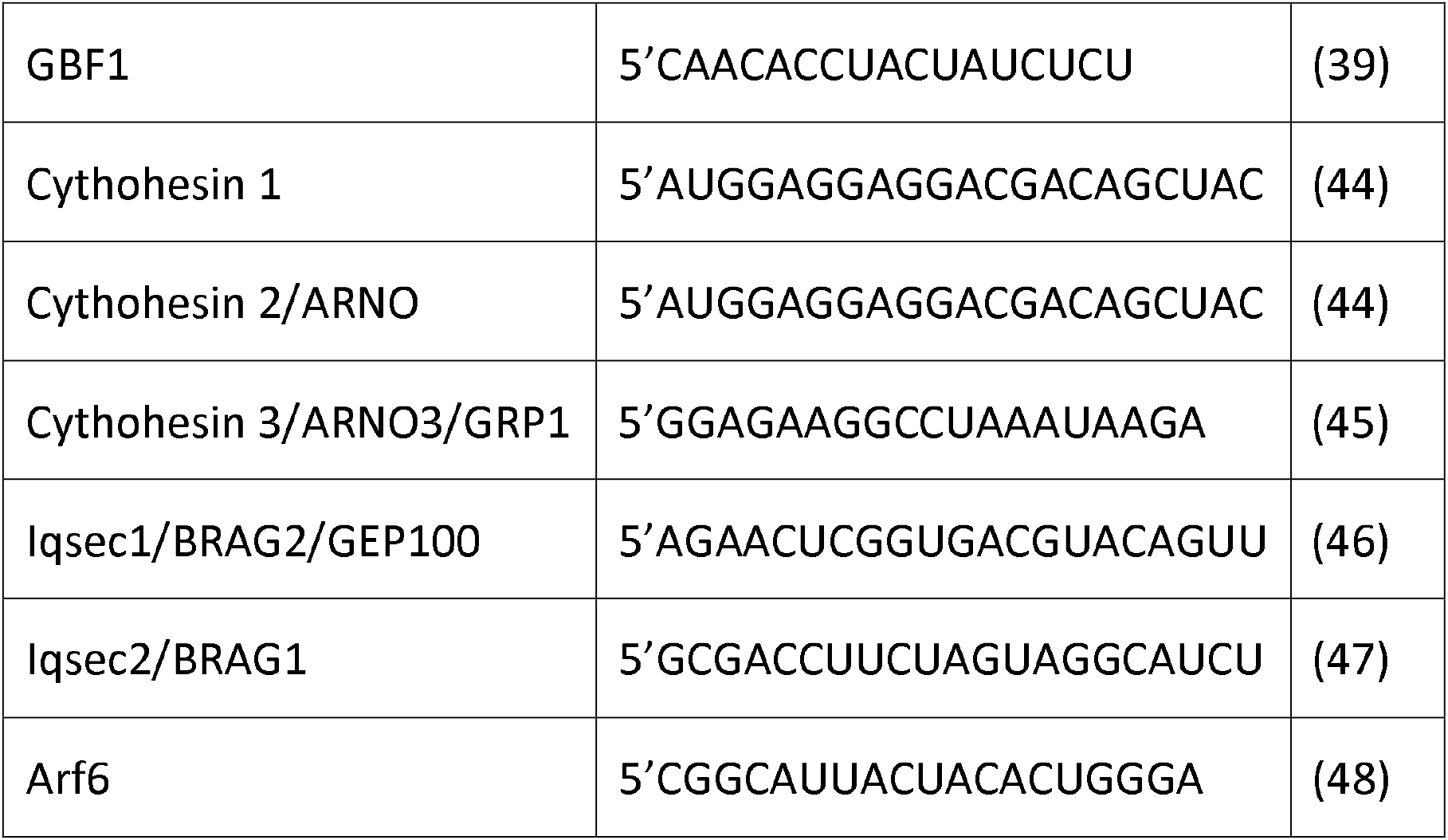

SiRNA duplexes were synthesized by Dharmacon with 3’UU overhangs, except for siRNA pool targeting Psd3/EFA6D which was purchased from Santa Cruz Biotechnology (sc-77475). Non-targeting siRNA control was from Ambion (siControl 1). siRNAs were transfected into HeLa cells grown on 96 well plates using Dharmafect-1 reagent (Dharmacon) according to the manufacturer’s recommendations. The replication experiments were performed ∼72h post siRNA transfection except for cells treated with siRNA targeting GBF1 which were first assessed at 48h post-transfection because prolonged depletion of GBF1 induces significant cytotoxicity. Cell viability was determined with CellTiter-Glo assay (Promega).

### Replication assay

Replicon RNA preparation and replicon assays were performed essentially as described in (42). Briefly, cells grown on white 96-well plates with clear bottoms were transfected with a polio replicon RNA expressing the Renilla luciferase gene in the complete medium supplemented with 30 μM of cell-permeable Renilla luciferase substrate Enduren (Promega). The plates were sealed with a transparent oversheet and incubated directly in the heated chamber of a multifunctional M3 microplate reader (Molecular Devices), the luciferase signals were measured every hour for 18 h post replicon transfection. The total replication signal was calculated as an area under the curve of kinetic measurement from 1 to 8 hours of replicon replication. The signal for each sample was averaged from at least 12 wells.

### Arf-EGFP pull-down assays

HeLa cells expressing Arf1-Venus fusion protein (33) grown on a six-well plate were infected at an MOI of 10 with the indicated viruses. For co-IP of individually expressed 2C and Arf1, HeLa cells were co-transfected on a six-well plate with a 3:1 mass ratio mix of pM1-2C and pArf1-EGFP plasmids (or a plasmid expressing EGFP instead of Arf-EGFP in controls) using Mirus 2020 transfection reagent (Mirus Bio). Infected cells were processed at 6, 8, and 18 h p.i. for poliovirus, Coxsackie virus B3, and rhinovirus 1A, respectively; those transfected with plasmids were incubated overnight (∼18h). The cells were either directly lysed in 750 μl of the mild lysis buffer (0.1M Tris-HCl pH 7.8 with 0.5% Triton X100 supplemented with a protease inhibitor cocktail (Millipore-Sigma)), followed by clearance of the lysate by low-speed centrifugation, or first processed for fractionation with a Cell Fractionation Kit (Cell Signaling, Cat.# 9038) according to the manufacturer’s protocol for cytoplasmic and total membrane and nuclear fractions. The proteins from the total membrane and nuclar fraction were similarly extracted with the mild lysis buffer. The cleared mild lysis buffer lysates were further used for pull-down assays.

The lysate protein concentration was determined using Bradford reagent (Bio-Rad), and Co-IP was performed using GFP Selector resin (Nanotag Biotechnology) according to the manufacturer’s protocol. Briefly, the amount of lysate containing 1 mg of total protein was mixed with pre-washed GFP Selector resin and incubated with rotation for 1 hour at 4°C. The beads were collected by low-speed centrifugation, and the unbound proteins were removed by extensive washing of the beads three times in the mild lysis buffer. The bound proteins were eluted by incubating the collected beads in a 2X denaturing Laemmli sample buffer at 95°C for 5 minutes.

### Microscopy and image processing

Cells were grown on glass coverslips in 12-well plates and fixed with 4% formaldehyde (Electron Microscopy Sciences) in PBS. Cells expressing Arf-Venus fusions were imaged with a confocal Zeiss LSM 510 microscope without any further processing to preserve the intracellular membranes. For immunofluorescent assays, the cells were permeabilized with 0.2% Triton X100 in PBS for 5 min, washed 3 times with PBS, and sequentially incubated for one hour with primary and secondary antibodies diluted in PBS with 3% blocking reagent (Amersham) with 3x PBS washes after each antibody incubation. The Structural Illuminated Microscopy (SIM) superresolution images were taken with a Nikon A1R microscope. Digital images were converted to TIFF format by the corresponding microscope software and processed with Adobe Photoshop for illustrations. All changes were applied to the whole image and images from the same experiment were processed with the same software settings.

### Statistical analysis

Data processing and statistical calculations were performed using the GraphPad Prism 9 package. Graphs show average values with standard deviation bars, the difference between pairs of experimental values was evaluated with unpaired (two-sample) t-test, p-values ≤0.05 were considered statistically significant. P-values ≤0.05 are designated by *, those ≤ 0.01 with **, those ≤ 0.001 with ***, and those ≤ 0.0001 with ****.

## Results

### Selection of poliovirus mutants resistant to high concentrations of BFA

The previously described poliovirus mutants capable of replicating in the presence of BFA were selected at 1-2 μg/ml of the drug. These viruses contained single amino-acid substitutions in the 2C and 3A proteins, but how these mutations restored viral replication in the presence of the drug could not be determined (37, 38). To get a better insight into the mechanism of BFA resistance, we set out to select viruses resistant to higher concentrations of the inhibitor to obtain more conspicuous phenotypes. Treatment of cells with BFA induces a dose-dependent inhibition of the secretory pathway and of Golgi-associated enzymatic activities such as protein glycosylation. The working concentrations of BFA up to 100 μg/ml (∼350 μM) have been used in different systems and the effects were reported to be fully reversible upon the drug removal (49-53).

First, we tried to select resistant variants by using high initial concentrations of BFA. HeLa cells were infected with wt poliovirus type I Mahoney at 50 PFU/cell and incubated in the presence of 20 or 50 μg/ml BFA for 6 hours (the duration of the poliovirus replication cycle in HeLa cells). Half of the total virus yield was used for the next passage in the same conditions, and the process was repeated for 10 passages. The cells tolerated six-hour incubation in the presence of the drug without any obvious signs of cytotoxicity, and previously such a selection scheme at 2 μg/ml of BFA resulted in a rapid establishment of a resistant viral population (38). However, after 10 passages at either 20 μg/ml or 50 μg/ml of BFA the resistant variants did not emerge. To facilitate selection, we first performed 10 passages at 4 μg/ml of BFA. The resistant population was successfully established, and sequencing of the 2C3A region of five plaque-purified viruses revealed that all of them shared a mutation C4200T (2C/S26L) located in the amphipathic helix of 2C (here and further on, mutations are designated by their position in the nucleotide sequence in the poliovirus genome and, in parenthesis, by the amino-acid position in the corresponding viral protein). This mutation was previously described in a BFA-resistant poliovirus mutant (38). Two of the isolates contained an additional 2C mutation A4286C (2C/M78L), two others shared another mutation T4410C (2C/V96A), and one of them had an additional mutation A5067G (2C/N135S). Three isolates shared a 3A mutation G5211A (3A/R34K), and the other two had different mutations in 3A, A5223G (3A/Q38G) or T5238C (3A/I43T), respectively (Figure S1).

The material from passage 10 at 4μg/ml BFA was used for further passaging 10 times at 20μg/ml of the drug. This time the selection was successful, and we sequenced 2C3A regions of six plaque-purified isolates, as well as that of the total viral RNA isolated from the material from passage 10. All of the individual isolates shared three 2C mutations: C4200T (2C/S26L), A4286C (2C/M78L), and A4318C (2C/Q65H). One of them had an additional 2C mutation A4380G (2C/Q86R). All but one isolate shared a 3A mutation G5211A (3A/R34K), and one had a 3A mutation T5238C (3A/I43T) (Figure S1). The sequence of 2C3A region of the total viral RNA revealed that only the mutations C4200T (2C/S26L) and A4318C (2C/Q65H) were strongly fixed in the population. Only the mutant nucleotide T was detected in position 4200, while in position 4318 there was a mix of either C or T, both of which would result in the same amino-acid change 2C/Q65H. All other mutation sites demonstrated a mixture of the wt and mutant sequences. We also detected a mixed sequence at the position of G4361A (2CV80I), a mutation that was described previously to be associated with BFA resistance (37) (Figure S1). Thus, the population of viruses propagated at 20μ/ml of BFA was still heterogeneous, and the number of mutations in 2C was increasing.

To see if the resistant population could resolve to the fittest genotype, the material from passage 10 at 20μg/ml of BFA was used for further passaging 25 times at 50 μg/ml of the drug. To avoid the possibility of missing BFA-dependent viruses, this time we did not plaque-purify individual isolates. Instead, a PCR fragment encompassing the 2C3A coding region of the poliovirus genome obtained from the total RNA was ligated to a plasmid vector, and 13 individual clones, as well as the original PCR material, were sequenced. All of the individual clones had a single 3A mutation T5238C (3A/I43T). Eleven of them had the same five mutations in 2C, C4200T (2C/S26L), A4318C (2C/Q65H), C4339A (2C/H72Q), T4410C (2C/V96A) and A4542G (2C/N140S). One clone did not have the A4542G (2C/N140S) mutation but instead contained another 2C substitution G4703A (V194I), and one clone had an additional 2C mutation C4851T (2C/A243V). Analysis of the sequence of the total PCR material revealed that the population selected at 50 μg/ml was much more homogenous than that after selection at 20 μg/ml of BFA, however in mutated positions 4410 and 4542 in 2C and 5238 in 3A, at least traces of the wt sequence could still be detected (Figure S1).

The three different 2C3A genotypes (clones 46, 59, and 60) found in the population resistant to 50 μg/ml were reconstructed in a replicon RNA coding for the *Renilla* luciferase gene instead of the polio capsid proteins (Fig. 1A). A comparison of replication of these three replicons at 25 and 50 μg/ml of BFA demonstrated that genotype 59 (the most abundant among the individual PCR fragments sequenced) was replicating better than the others (Fig. 1B), and it was selected for further analysis. This mutant replicated in the broad range of BFA concentrations, and showed a mild BFA-dependent enhancement of replication in HeLa cells (Fig. 1C).

**Figure 1.**
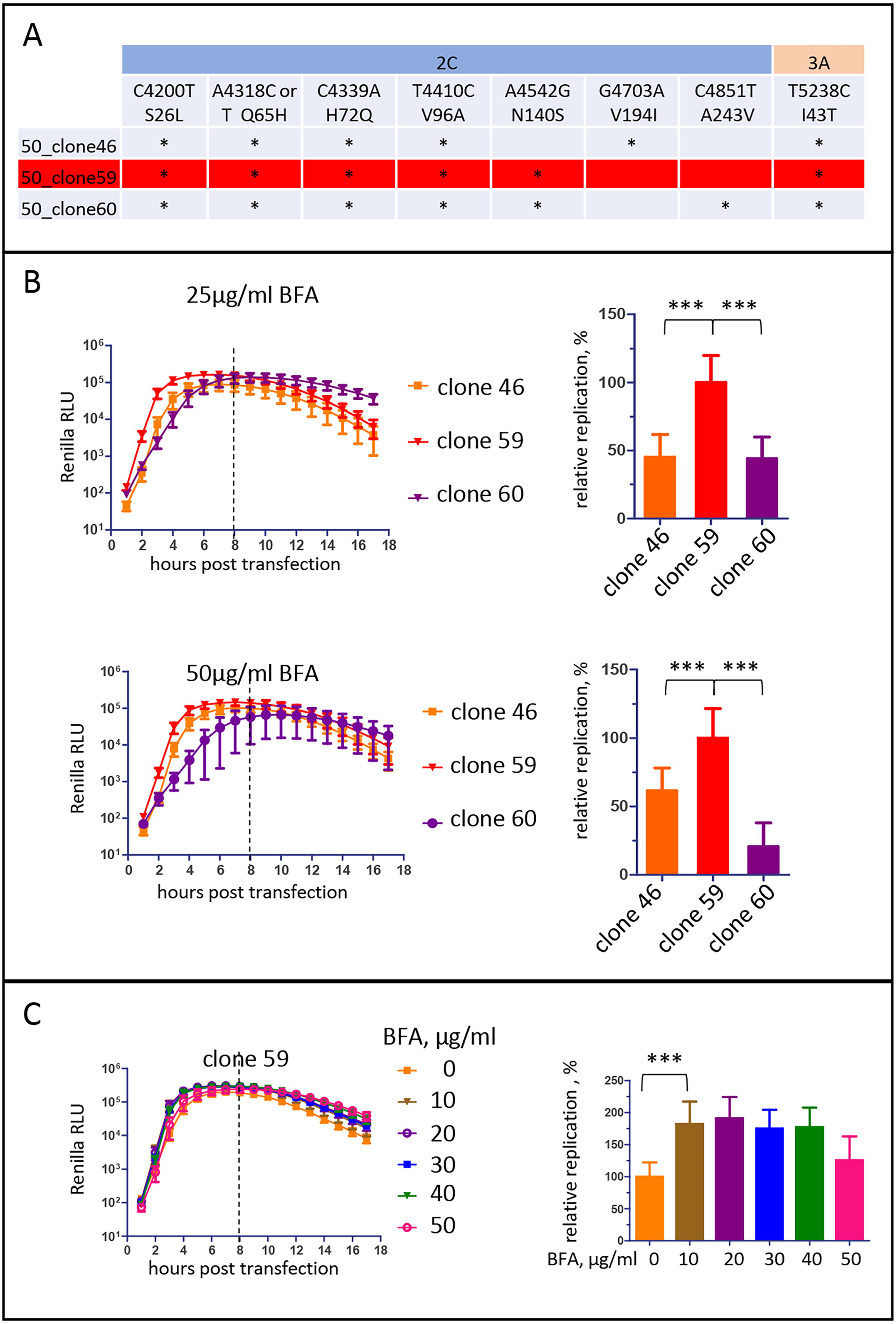
Selection poliovirus mutants resistant to high BFA concentration. **A**. A summary of mutations in 2C and 3A proteins found in highly BFA-resistant poliovirus isolates. **B**. Replication of Renilla replicons with mutations found in highly resistant genotypes at the indicated concentrations of BFA. Relative replication is calculated for the signal up to 8 h post-transfection (dashed line) and is normalized to the replication of the mutant 59.

Thus, poliovirus resistance to high concentrations of BFA requires the accumulation of multiple mutations in 2C, and can be achieved only upon the sequential increasing concentration of the drug.

### Mutations in 2C are responsible for the high-level resistance to BFA

The mutations in 2C were concentrated in the N-terminal part of the protein which contains an amphipathic helix and is important for the recruitment of 2C to membranes (54, 55). The mutation in 3A was localized in the stretch of amino-acids where other mutations conferring resistance to BFA and the inhibitors of the OSBP-PI4KIIIβ-dependent enrichment of the replication organelles in PI4P and cholesterol were reported (37, 38, 56) (Fig. 2A). To understand the contribution of individual mutations, we first constructed replicons containing only 2C or 3A mutations from clone 59 (59/2C and 59/3A, respectively) and assessed their replication at low (2 μg/ml) and high (25 μg/ml) concentrations of BFA. Replicon 59/3A with only the T5238C (3A/I43T) mutation replicated much better without BFA than the parental replicon 59, but was severely compromised in the presence of either concentration of the inhibitor (Fig. 2B). Replicon 2C/59 also replicated better without the drug than the parental replicon 59, although not as robustly as the replicon 59/3A. Its replication was resistant to either high or low concentrations of the drug but was not stimulated by BFA like that of replicon 59 (Fig. 2B). In different experiments, the total replication signal of 2C/59 at the high concentration of BFA sometimes demonstrated minimal but statistically significant decrease compared to the replication without the inhibitor. Thus, the five 2C mutations are sufficient to confer the high level of BFA resistance but not the BFA-dependent phenotype in HeLa cells.

**Figure 2.**
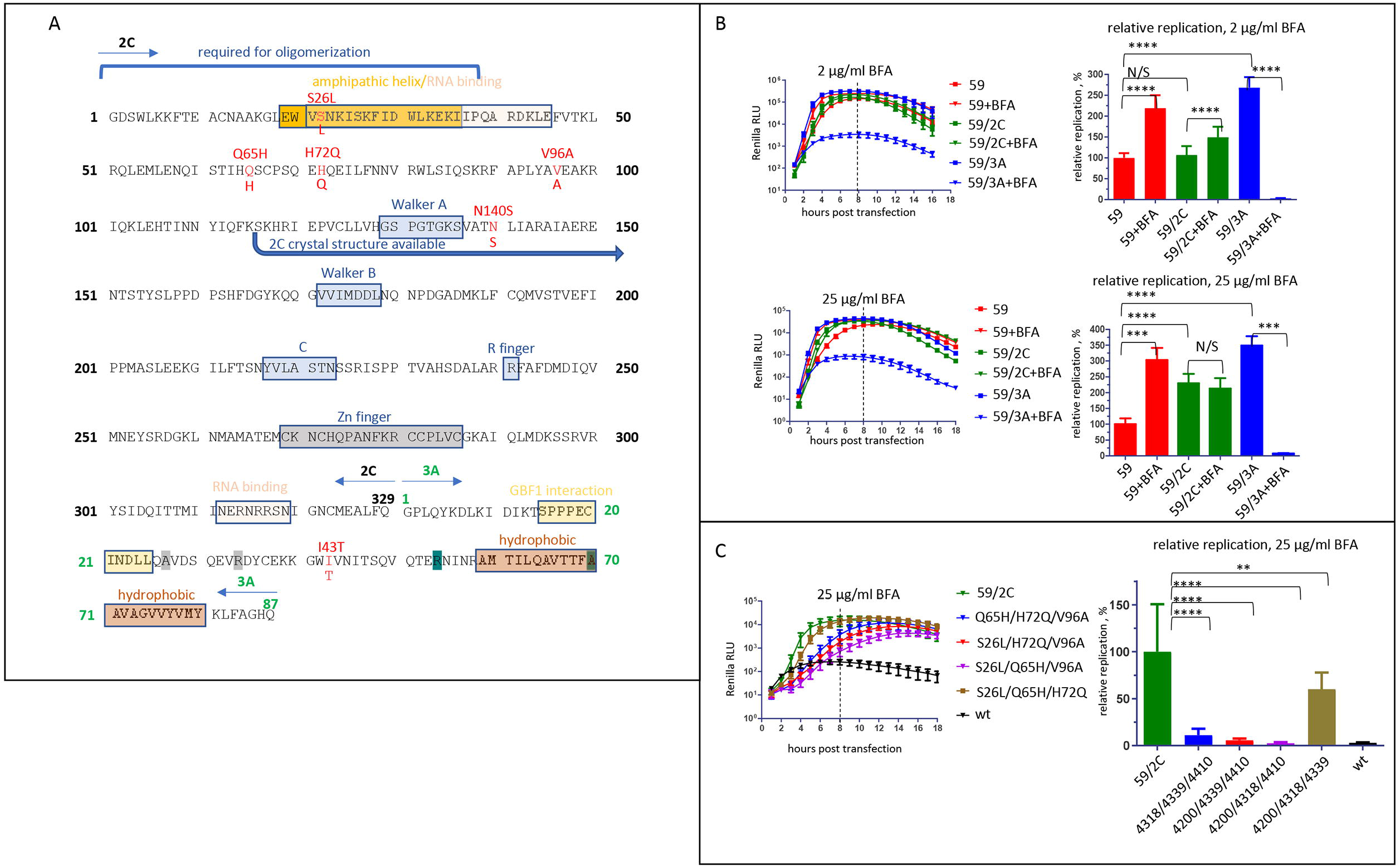
Mutations in 2C are necessary and sufficient to impart high level of BFA resistance. **A**. A schematic of the 2C-3A poliovirus polyprotein fragment with black and green numbers indicating 2C and 3A amino-acids, respectively. Known functional motives are designated by colored boxes, and BFA-resistant mutations of genotype 59 are in red. The grey and teal highlights of the 3A amino-acids indicate locations of previously reported mutations conferring resistance to BFA and enviroxime-like compounds, respectively. **B**. Replication of Renilla replicons bearing the full complement of mutations from genotype 59, or those in the 2C or 3A proteins only. Relative replication is calculated for the signal up to 8 h post-transfection (dashed line) and is normalized to that of 59. **C**. Replication of Renilla replicons with the indicated combinations of mutations in 2C from the genotype 59. Relative replication is calculated for the signal up to 8 h post-transfection (dashed line) and is normalized to that of 59/2C.

To see which 2C mutations contributed the most to the highly resistant phenotype, we first constructed replicons lacking a single mutation out of the five present in 59/2C. The removal of the most N-terminal mutation C4200T (2C/S26L) did not affect the replication at 25μg/ml of BFA, while the replicons lacking A4318C (2C/Q65H) and C4339A (2C/H72Q) mutations replicated at about half the efficiency of the control 59/2C replicon (Figure S2). Interestingly, the removal of the C-terminal mutations T4410C (2C/V96A) or A4542G (2C/N140S) noticeably increased the level of replication in the presence of 25 μg/ml of BFA (Figure S2). We then generated mutants containing combinations of 3 mutations in the background of a replicon lacking A4542G (2C/N140S) (the most C-terminal mutation that was dispensable for BFA-resistant phenotype). In the presence of 25 μg/ml of BFA only the replicon containing the three most N-terminal mutations C4200T (2C/S26L), A4318C (2C/Q65H), and C4339A (2C/H72Q) replicated similar to the control 59/2C replicon, although with a delayed kinetics. Replication of all other replicons was severely compromised, even though they still showed BFA resistance compared to the wt replicon (Fig. 2C).

Thus, the BFA resistance can be conferred by multiple mutations in 2C in the context of wt 3A, and the combination of N-terminal mutations C4200T (2C/S26L), A4318C (2C/Q65H) and C4339A (2C/H72Q) contributes the most to the resistant phenotype.

### The mutant virus demonstrates a strong BFA-dependent phenotype in a cell-type specific manner

We analyzed the BFA-resistant phenotype of the replicon 59 in cells of different origins. In all cell lines tested, the control wt replicon replicated efficiently and was strongly inhibited by BFA (Fig. 3A). Vero (green monkey kidney) was the only cell line where replicon 59 replicated to the levels comparable to that of the wt replicon in the presence and in the absence of the drug, i.e. there was no BFA-dependent phenotype (Fig. 3A). In all other cell lines, the level of the replication of the BFA-resistant replicon approached that of the control replicon only in the presence of the drug, while in the absence of the inhibitor, its replication varied significantly. In HeLa (human cervical carcinoma) and HEK293 (human embryonic kidney) cells, the total replication signal of the replicon 59 was within 25% of that of the control. In Huh7 (human hepatoma), MG63 (human osteosarcoma), and A549 (human lung carcinoma) cells the total replication signal of replicon 59 in the absence of BFA was less than 10% of the control wt replicon replication (Fig. 3A). In Caco2 (human colorectal adenocarcinoma) cells the defect of replicon 59 replication was particularly severe, where only a trace replication signal in the absence of the drug could be observed, while in the presence of BFA it was recovered to nearly the level achieved by the wt replicon without the inhibitor (Fig. 3A).

**Figure 3.**
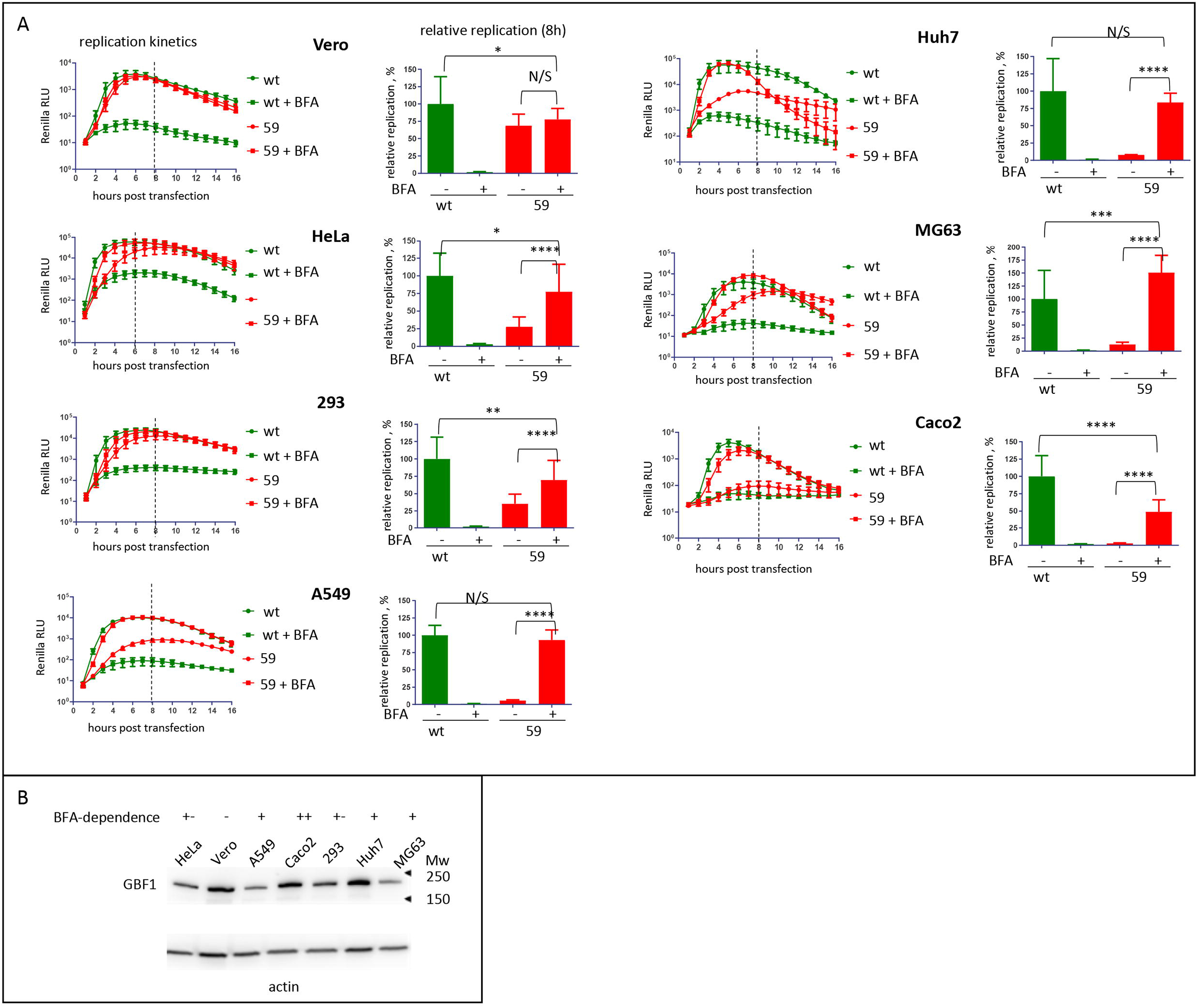
Replication of the BFA-resistant mutant is cell-type specific. **A**. Replication of wt replicon and the replicon with mutations in 2C and 3A from genotype 59 in the indicated cell lines in the absence or presence of 25 μg/ml of BFA. Relative replication is calculated for the signal up to 8 h post-transfection (dashed line) and is normalized to the replication of wt replicon without BFA. **B**. Expression of GBF1 in the cell lines used in A. Actin represents a loading control.

Previously, the analysis of the replication of a poliovirus mutant resistant to a low concentration of BFA which contained single amino-acid substitutions in the 2C and 3A proteins showed that it was dependent on the expression of GBF1 (38, 39). Thus, we compared the level of GBF1 expression in different cell lines. No correlation between the levels of GBF1 and the manifestation of the BFA-dependent phenotype of replicon 59 could be observed (Fig. 3B). These data show that high level of BFA resistance imparts a significant fitness cost in a cell-type-specific manner and demonstrates the importance of cell factors for the functioning of the viral replication machinery.

### Replication of the highly BFA-resistant mutant is independent of GBF1 levels

We investigated the effect of GBF1 depletion on the replication of the highly BFA-resistant mutant in HeLa cells. At 48h post siRNA treatment, the amount of GBF1 was significantly decreased (Fig. 4A, GBF1 western blot). Surprisingly, the replication of replicon 59 was stimulated in GBF1-depleted cells in the absence of BFA, while the replication in the presence of the drug was similar in cells treated with either control or anti-GBF1 siRNA (Fig. 4A, 48h). In cells incubated with siRNA for 72h, further reduction of GBF1 expression was observed (Fig. 4A, GBF1 western blot), but at this time point cells treated with GBF1 siRNA showed noticeable toxicity. In cells treated with control siRNA for 72h, the BFA-dependence of replication of mutant 59 became even more pronounced than at 48h, increasing up to ∼25x compared to the replication without the drug (Fig. 4A, 72h). In GBF1-deleted cells, the replication was similar in the presence and in the absence of BFA, and in both conditions the replicon replicated better than in control cells without the drug (Fig. 4B, 72h). Thus, the reduction of GBF1 levels improved viral replication in the absence of BFA, and did not affect BFA resistance.

**Figure 4.**
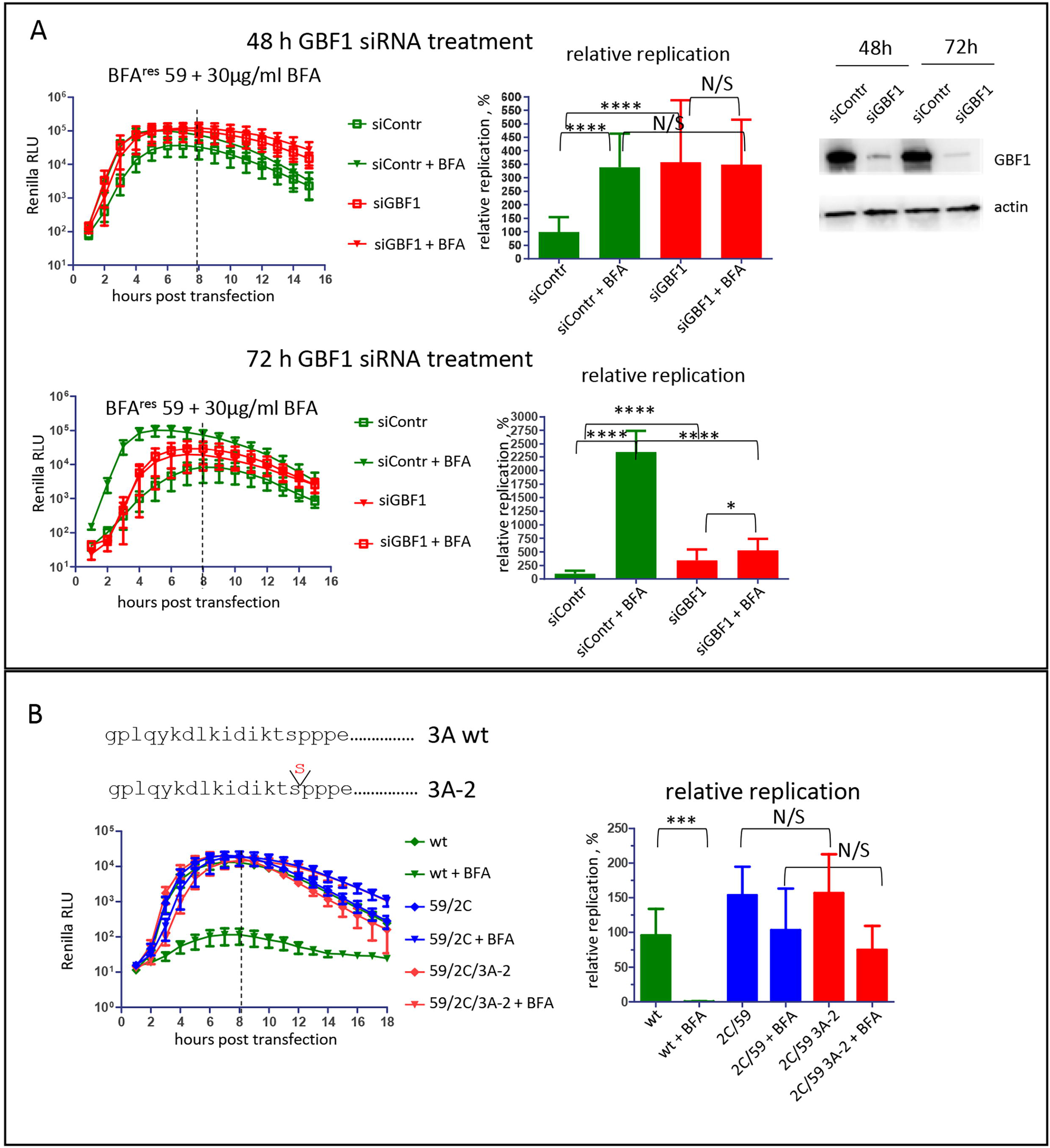
BFA-resistance conferred by mutations in 2C is independent of GBF1 levels. **A**. Replication of the Renilla replicon with mutations from genotype 59 in cells treated with siRNA against GBF1 for 48 (top) and 72 h (bottom) with or without 30 μg/ml BFA. Western blot shows the level of GBF1 depletion at the time of replicon transfection. Actin is represents a loading control. Relative replication is calculated for the signal up to 8 h post-transfection (dashed line) and is normalized to the replication in cells treated with control siRNA without BFA. **B**. Replication of Renilla polio replicons (wt, with mutations in 2C form the genotype 59 (2C/59), and with a combination of BFA resistant mutations in 2C and mutation 3A-2 in 3A that inhibits the 3A-GBF1 interaction (2C/59/3A-2)) with or without 25 μg/ml BFA. Relative replication is calculated for the signal up to 8 h post-transfection (dashed line) and is normalized to the replication of the wt replicon without BFA.

To confirm that the 2C-mediated mechanism of high level BFA resistance does not depend on GBF1, we introduced the 2C mutations in the context of the so-called 3A-2 mutation, insertion of an additional serine at the 13^th^ position of the protein 3A (Fig. 4B). The 3A-2 mutation in poliovirus 3A or analogous mutation in 3A of Coxsackie virus B3 severely impairs the 3A-GBF1 interaction and reduces GBF1 recruitment to replication organelles (12, 34). Accordingly, we previously demonstrated that the 3A-2 mutation significantly increases the sensitivity of poliovirus replication to BFA and compromises the efficacy of BFA-resistant mutations in 2C or 3A selected at low concentrations of the drug (36, 38). However, we observed that the replicon 2C/59/3A-2 with the 2C of the highly resistant virus in the context of 3A-2 replicated in the presence and in the absence of BFA similar to the control 2C/59 replicon containing the wt 3A sequence. This further supports that GBF1 is unlikely to play a major role in the BFA-resistant phenotype of this mutant (Fig. 4B).

Collectively, these data strongly suggest that the mechanism of BFA resistance conferred by mutations in 2C may be GBF1-independent.

### Replication of the mutant virus induces Arf activation even in the presence of BFA

We recently demonstrated that enterovirus replication is accompanied by the simultaneous recruitment of all cellular Arfs to the replication organelles (33). To monitor Arf recruitment, we infected HeLa cell lines expressing individual Arf-Venus fusions with an MOI of 50 of the mutant 59 in the absence and in the presence of 25 μg/ml of BFA. The cells were fixed at 6 h p.i and imaged without additional immunostaining to maximally preserve intracellular membranes. In mock-infected cells, Arfs 1, 3, 4 and 5 were released from the intracellular membranous structures in the presence of BFA. As expected, Arf6, which is activated by BFA-insensitive ArfGEFs and controls membrane trafficking associated with the plasma membrane was not significantly affected (Fig. 5A, mock panel). However, in cells infected with the mutant 59, we observed strong recruitment of all Arfs to the perinuclear replication organelles irrespective of the presence of BFA (Fig. 5A, PV panel).

**Figure 5.**
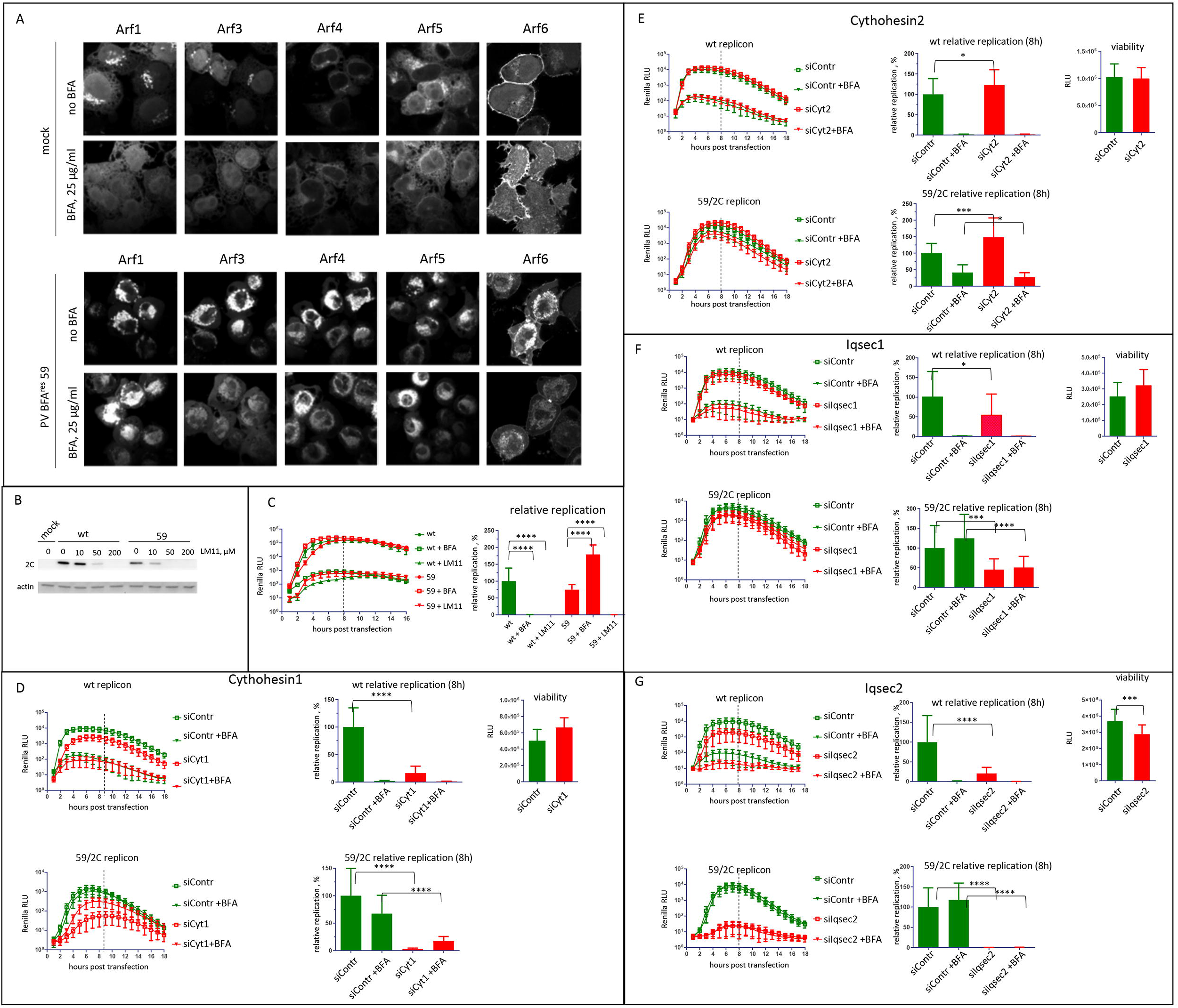
Arfs are recruited to the replication organelles of the BFA-resistant mutant even in the presence of BFA. **A**. HeLa cell lines stably expressing individual Arf-Venus fusions were infected (or mock-infected) with the BFA-resistant mutant 59 at an MOI of 10 and incubated with or without 25 μg/ml of BFA for 6 h. **B**. HeLa cells were infected with wt poliovirus or the BFA-resistant mutant 59 at an MOI of 10 and incubated with the indicated concentrations of the pan-ArfGEF inhibitor LM11 for 6 h. Accumulation of the viral protein 2C reflects replication (actin represents a loading control). **C**. Replication of Renilla polio replicons (wt and the BFA-resistant genotype 59) in the presence of 25 μg/ml BFA or 200 μM of LM11. Relative replication is calculated for the signal up to 8 h post-transfection (dashed line) and is normalized to the replication of wt replicon without the inhibitors. **D-G**. Replication of Renilla polio replicons (wt and the BFA-resistant 59/2C mutant) in HeLa cells treated with siRNA targeting individual ArfGEFs with or without 25 μg/ml of BFA. The replication assay was performed 72h post-siRNA transfection, viability data reflect the cell state at the end of the replication assay. Relative replication is calculated for the signal up to 8 h post-transfection (dashed line) and is normalized to the replication of each replicon in cells treated with control siRNA without the inhibitor.

To determine if Arf activation is required for the replication of the resistant mutant, we used LM11, a compound that inhibits the activity of both, the BFA-sensitive and the BFA-insensitive ArfGEFs *in vitro* and *in vivo* (32). Replication of wt virus and mutant 59 were inhibited by LM11 in a dose-dependent manner (Fig. 5B). Importantly, cells incubated with LM11 during the infection did not show any signs of cytotoxicity. The inhibitory effect of LM11 was confirmed in a replicon replication experiment. The replication of the wt replicon was similarly inhibited by both BFA and LM11, while replication of the replicon 59 showed strong BFA resistance as expected, but was inhibited by LM11 similar to the wt replicon (Fig. 5C). Thus, the BFA-resistant virus still requires Arf activation for replication.

Given that Arf activation is still observed in cells infected with the mutant virus when BFA-sensitive GEFs (such as GBF1) are inhibited implies Arf activation by BFA-insensitive ArfGEF(s). The human genome encodes 14 proteins with functional Sec7 domains, only three of which (GBF1, BIG1 and BIG2) are BFA-sensitive. The Sec7 domains of different ArfGEFs can likely activate all Arfs, and the specificity of Arf activation by different ArfGEFs in cells is determined by poorly understood targeting signals restricting specific Arfs and ArfGEFs to a particular subcellular location (reviewed in (30)). The proteomics and transcriptomics data show that HeLa cells express several BFA-insensitive ArfGEFs: Cythohesin1, Cytohesin2/ARNO, Cytohesin3/ARNO3/GRP1, Iqsec1/BRAG2/GEP100, Iqsec2/BRAG, and Psd3/EFA6D (57, 58). We first screened if siRNA-mediated depletion of these BFA-insensitive ArfGEFs affects the replication of 2C/59 replicons in HeLa cells in the presence of 25μg/ml BFA. Depletion of PSD3 and Cytohesin3 had no effect on replication (Fig. S3). Depletion of Cytohesin1 or Cytohesin2 inhibited the replication to 30-50% of control, while the depletion of Iqsec1 and especially Iqsec2 had a much more pronounced inhibitory effect (Fig S3). We further analyzed the effect of depleting Cytohesin1, 2, and Iqsec1, 2 on the replication of both 59/2C and wt replicons in the presence and absence of 25μg/ml BFA. In cells treated with siRNA targeting Cytohesin1, the replication of both replicons without BFA was significantly inhibited, but the BFA-resistant phenotype of 59/2C was still observed (Fig. 5, D). The depletion of Cytohesin2 and Iqsec 1 had minimal effect on the replication of either of the replicons with and without BFA (Fig 5. E, F). The depletion of Iqsec2 inhibited replication of the wt replicon without BFA to ∼25% of that in cells treated with control siRNA, and demonstrated the strongest suppression of the replication of the 59/2C mutant, both in the presence and absence of BFA, but the depletion of this ArfGEF was also noticeably toxic to cells (Fig. 5G).

Collectively, these results demonstrate that the 2C-mediated mechanism of high level BFA resistance could bypass the requirement for GBF1 activity, but still requires activated Arfs that are likely supplied by multiple BFA-insensitive ArfGEFs.

### Arf1 and Arf6, but not other Arfs provide the major contribution to the replication of BFA-resistant viruses in the presence of BFA

To determine which of the five Arf isoforms present in HeLa cells is required to support the replication of the 59/2C replicon in the presence of BFA, we utilized cells with CRISPR-CAS9 knockouts of individual Arf 1, 3, 4 and 5 (40), and siRNA-mediated depletion of Arf6. In control cells, the 59/2C replicon demonstrated the expected resistance to BFA with the total replication signal of ∼75% and ∼65% in the absence or presence of 25μg/ml of BFA, respectively, compared to the replication of the wt replicon without the inhibitor (Fig. 6A, HeLa control). The knockout of Arf1 and depletion of Arf6, but not the knockout of other Arfs specifically inhibited the replication of 5/2C9 replicon in the presence of BFA, from 45% down to ∼14% with and without BFA for Arf1, and from ∼55 to ∼25% for Arf6, relative to the replication of the wt replicon in the absence of the drug (Fig. 6).

**Figure 6.**
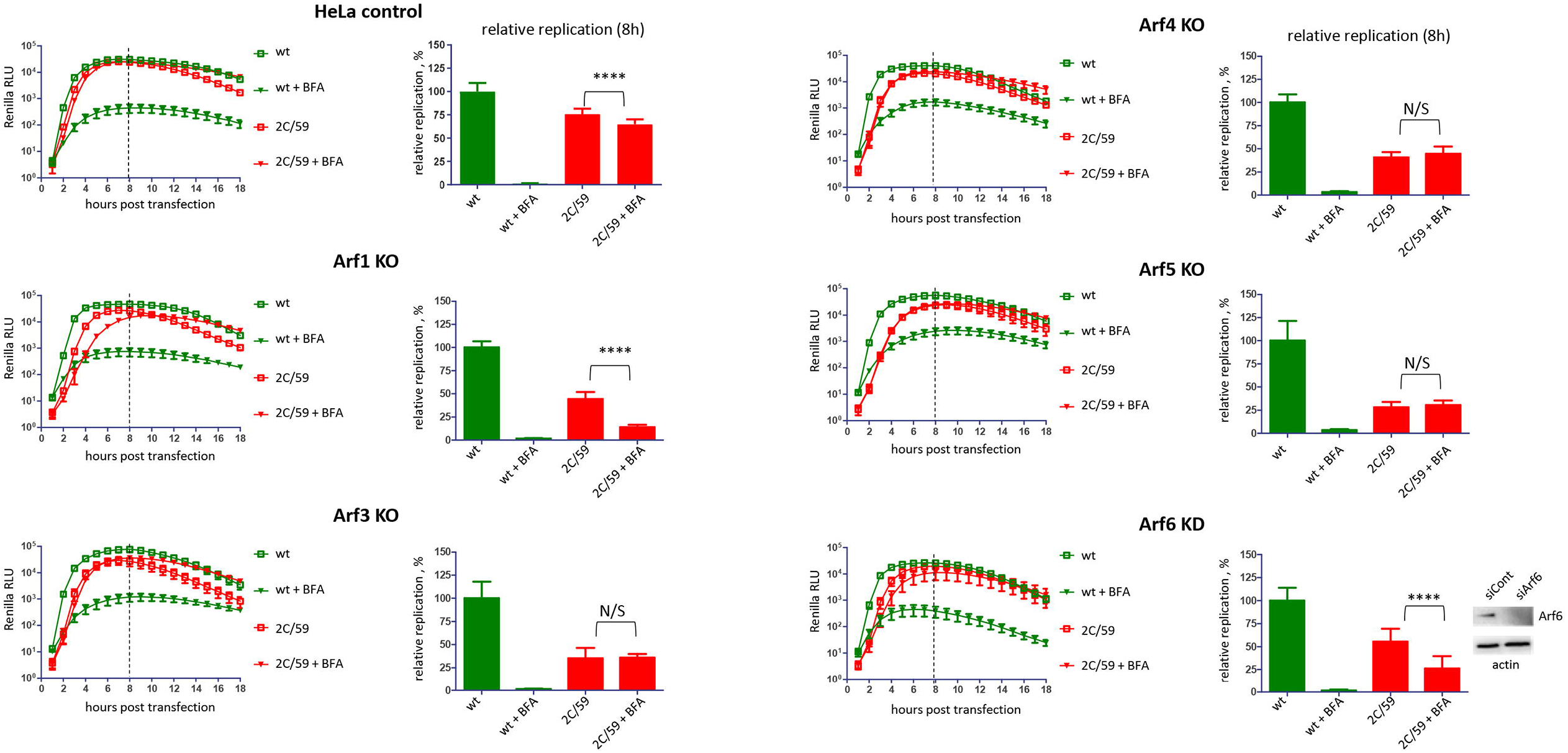
BFA-resistant phenotype conferred by mutations in 2C is sensitive to depletion of specific Arfs. Replication of the Renilla polio replicons (wt and the BFA-resistant mutant 59 (2C/59) in cells with CRISPR knockout of individual Arfs 1-5 and siRNA-knockdown of Arf6 with and without 25 μg/ml BFA. Western blot shows the level of Arf6 depletion at the time of replicon RNA transfection. Relative replication is calculated for the signal up to 8 h post-transfection (dashed line) and is normalized to the replication of wt replicon without BFA.

Thus, the BFA-resistant phenotype conferred by mutations in 2C still depends on Arfs, and Arf1 and Arf6 are the major contributors to viral replication in the presence of BFA.

### BFA-resistant mutations strongly increase the interaction of 2C with activated Arf1

The accumulation of Arfs on the replication organelles of wt and the BFA-resistant virus, in the latter case even in the presence of high concentrations of BFA, strongly suggests that activated Arfs perform an essential role in the replication process. Arfs function in cells by organizing biochemically distinct domains on membranes through direct interactions with Arf effector proteins. Given that Arf accumulation on membranes in the presence of BFA correlates with the accumulation of mutations in 2C, we hypothesized that 2C may engage in direct interaction with Arfs. To analyze 2C-Arf interactions, we co-expressed wt poliovirus 2C, or 2C from the resistant mutant 59 (2C/59) with individual Arf-EGFP fusions and performed co-IP with anti-GFP beads. In control samples, 2Cs were co-expressed with only EGFP. No specific interaction of the wt 2C with any of the Arfs could be detected in this assay, but the interaction of the mutant 2C/59 with Arf1, and to a lesser extent with Arf3, was markedly increased (Fig. 7A).

**Figure 7.**
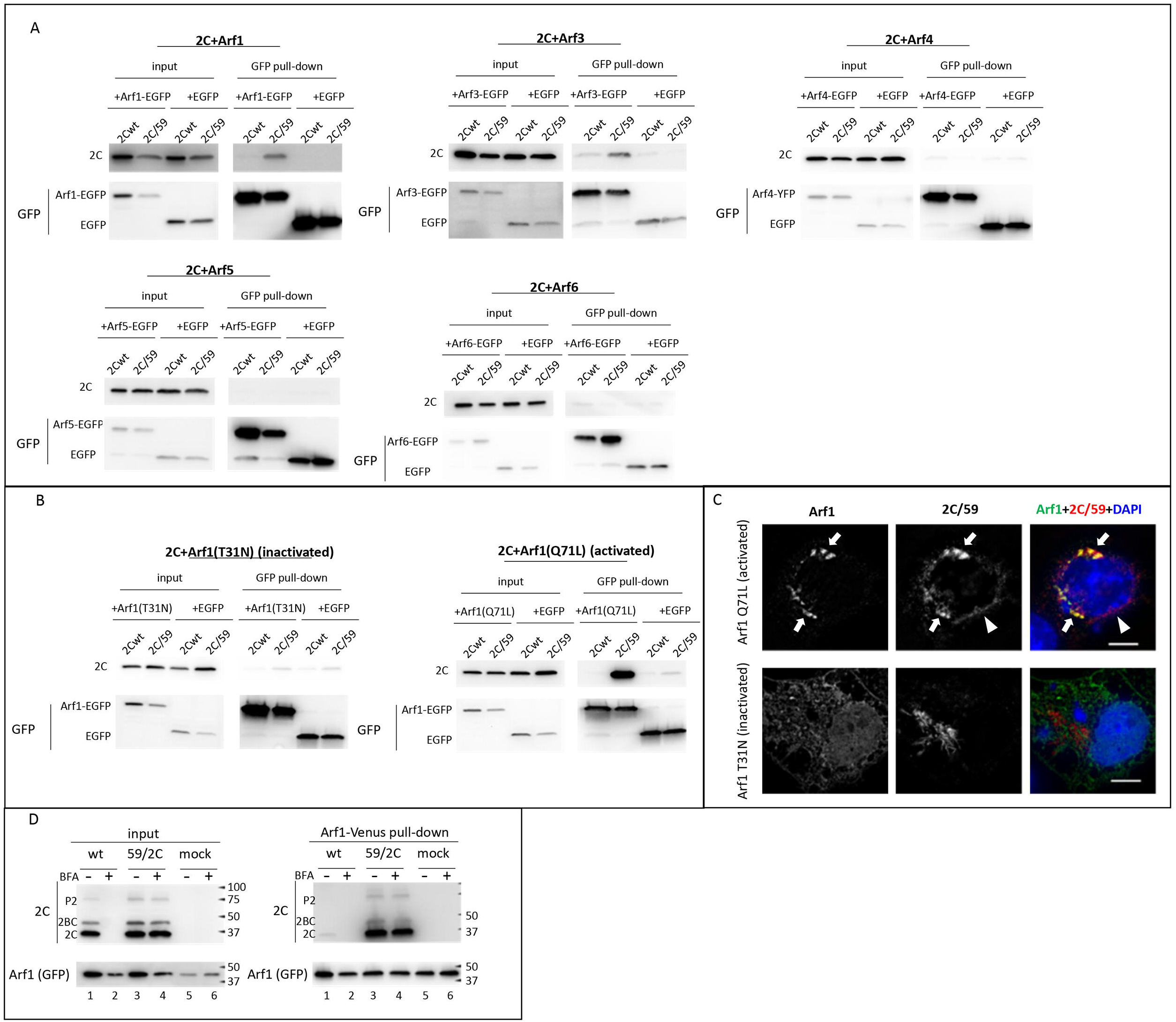
BFA-resistant mutations in 2C increase its interaction with activated Arf1. **A**. HeLa cells were cotransfected with plasmids expressing poliovirus 2C (wt or the BFA-resistant mutant (2C/59)) and those expressing individual Arf-EGFP fusions. In control cells, 2Cs were co-expressed with only EGFP. Pull-down was performed with anti-GFP beads and the amount of the recovered proteins was determined by western blot with anti-2C and anti-GFP antibodies. **B**. Hela cells were co-transfected with plasmids encoding 2Cs (wt or the BFA-resistant mutant (2C/59)) with EGFP fusions of Arf mutants T31N and Q71L locked in GDP (inactivated) or GTP (activated) states, respectivly. Pull-down was performed with anti-GFP beads and the amount of the recovered proteins was determined by western blot with anti-2C and anti-GFP antibodies. **C**. HeLa cells co-expressing 2C from the resistant mutant (2C/59) and EGFP fusions of activated (Q71L) or inactivated (T31N) Arf1 mutants were processed for immunofluorescence with anti-2C antibody and imaged by structural illumination superresolution microscopy. The scale bar is 10 μm. Co-localization of 2C/59 and Arf1/Q71L is indicated by arrows. Arrowhead marks 2C/59 signal separate from the Arf1/Q71L signal. **D**. HeLa cells stably expressing Arf1-Venus fusion were infected with an MOI of 50 of wt poliovirus or the BFA-resistant mutant (59/2C) and incubated with or without 25 μg/ml of BFA for 6 h. Cells were lysed and processed for membrane fraction separation which was used for co-IP with anti-GFP beads. The amount of recovered proteins was determined by western blotting with anti-2C and anti-GFP antibodies. Polyprotein fragments P2 and 2BC are marked.

We further focused on the interaction of 2C with Arf1 since this Arf1 contributes the most to the replication of both wt poliovirus and BFA-resistant mutant ((33) and data presented here). We performed similar co-IP experiments after co-expressing the wt and BFA-resistant 2Cs with Arf1/T31N locked in the inactive, GDP-bound conformation, and Arf1/Q71L locked in the activated, GTP-bound conformation (59). As above, we could not detect a specific interaction of wt 2C with any of the Arf mutants, but the 2C/59 mutant interacted very strongly with the activated Arf1/Q71L (Fig. 7B).

We also analyzed the cellular localization of 2C/59 relative to activated and inactivated Arf1 using structural illuminating microscopy (SIM) imaging. As can be seen in Fig. 7C, there was no co-localization of 2C/59 with inactive Arf1 T31N mutant, while in cells co-expressing 2C/59 and the activated Arf1 Q71L mutant the two proteins extensively co-localized (Fig. 7C, arrows). Still, there were also regions where only 2C/59 signal was present (Fig. 7C, arrowhead).

We further analyzed the 2C-Arf1 interaction under conditions of *bona fide* viral infection. HeLa cells stably expressing Arf1-Venus fusion were infected with an MOI of 50 of wt poliovirus, or the BFA-resistant mutant 59/2C with or without 25μg/ml of BFA, and the cells were processed for co-IP at 6 h p.i. We used the BFA-resistant mutant with mutations only in 2C because it replicates similarly in the presence and in the absence of BFA, in contrast to the mutant with mutations in both 2C and 3A which shows a BFA-dependent phenotype. Arfs are abundant cellular proteins, and only a fraction of the total Arf pool exists in the activated, membrane-bound form. Thus, to increase the signal of the activated, membrane-bound Arf1-Venus, which could be interacting with 2C, the co-IP was performed with the membrane fraction of the cellular lysates. The analysis of the input showed that the replication of the wt virus in the absence of the drug was similar to the replication of the 59/2C mutant in both conditions, and that BFA effectively blocked the replication of the wt virus, as expected (Fig. 7D, input, compare lanes 1 and 3, 4)). In the co-IP samples, we observed a trace signal of wt 2C (Fig. 7D, input, lane 1), but very strong signals of mutant 2C in the pull-down from infected cells incubated with or without BFA (Fig. 7D, pull-down, lanes 3, 4). Other mutant 2C-containing fragments of polyprotein processing (2BC and P2) were also recovered in Arf1 co-IP (Fig. 7D, pull-down, lanes 3, 4).

These data show that the BFA-resistant mutations in poliovirus 2C strongly increase its interaction with activated Arf1, suggesting a mechanism of BFA resistance based on the scavenging of activated Arfs generated by BFA-insensitive ArfGEFs.

### 2C-Arf1 interaction correlates with the sensitivity of enterovirus replication to BFA

We further analyzed the 2C-Arf1 interaction of diverse enteroviruses. Control HeLa cells and HeLa cells expressing Arf1-Venus fusion were infected with an MOI of 50 of poliovirus (a representative of Enterovirus C species), Coxsackie virus B3 (Enterovirus B species), or rhinovirus 1A (Rhinovirus A species). Since these viruses replicate with different kinetics, the co-IP from membranous fractions of the cellular lysates with anti-GFP beads was performed at 6 h p.i. for poliovirus, 8 h p.i. for Coxsackie virus B3, and 18 h p.i. for rhinovirus 1A. In agreement with the data shown above, there was no specific recovery of poliovirus 2C in co-IP with Arf1 Fig. 8A). Similarly, we did not observe a co-IP signal for 2C of Coxsackie virus B3 (Fig. 8B), but the interaction of 2C of rhinovirus 1A with Arf1 was very strong (Fig. 8C).

**Figure 8.**
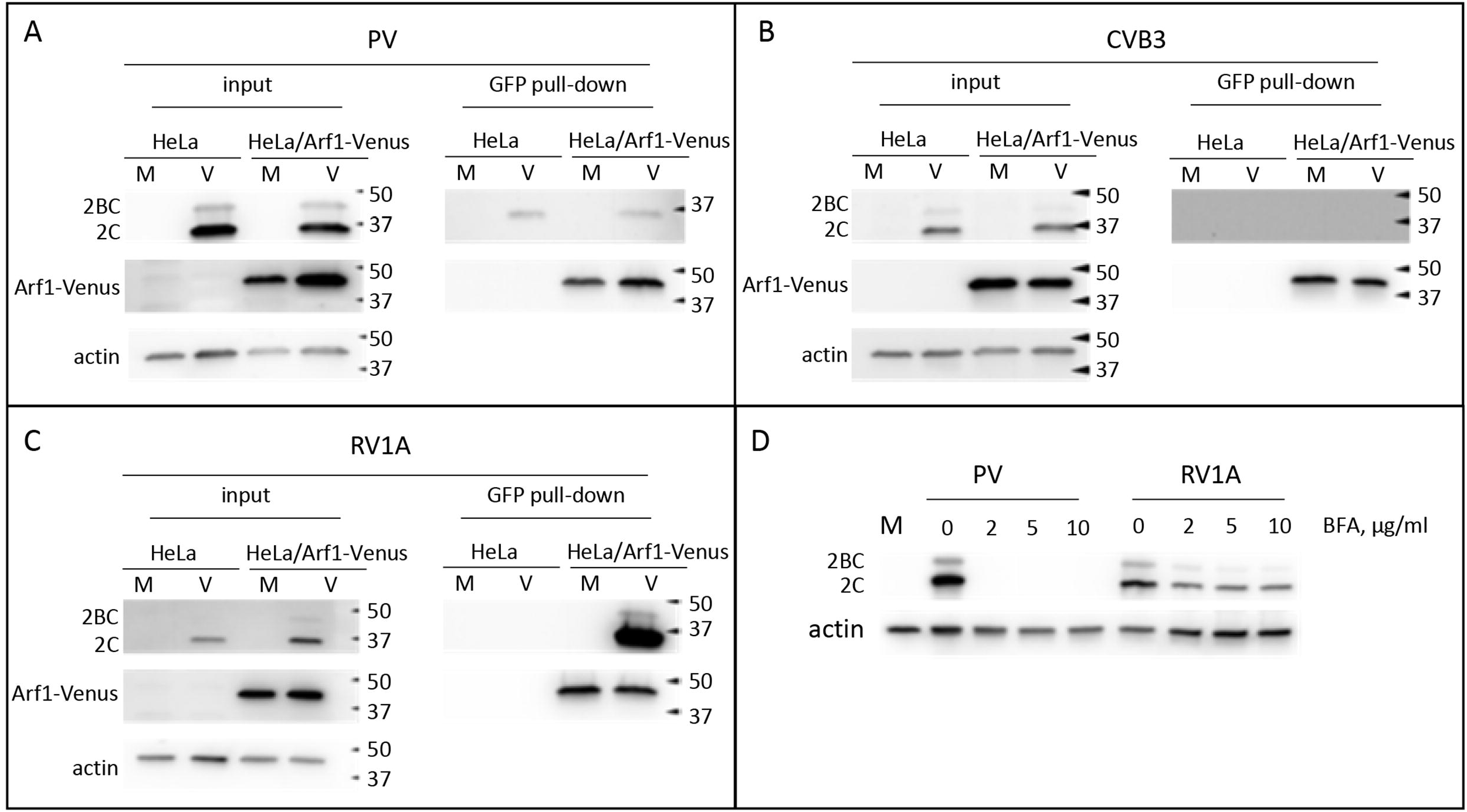
2C-Arf1 interactions correlates with BFA sensitivity of enterovirus replication. **A-C**. Control HeLa cells or HeLa cell stably expressing Arf1-Venus fusion) were infected (or mock-infected) with an MOI of 50 of poliovirus (A), Coxsackie virus B3 (B) or Rhinovirus 1A (C). The cells were lysed and processed for membrane fraction separation at 6 h p.i. for poliovirus, 8 h p.i. for Coxsackievirus B3 and 18 h p.i. for Rhinovirus 1A-infected cells. The membrane fractrion was used for co-IP with anti-GFP beads and the amount of recovered proteins was determined by western blotting with anti-2C and anti-GFP antibodies. Actin represents a control for co-IP input. **D**. HeLa cells were infected (or mock-infected) with an MOI of 10 of poliovirus or Rhinovirus 1A and incubated in the presence of the indicated concentrations of BFA for 8 h. Cells were lysed and the lysates analyzed by western blotting with anti-2C and anti-actin (loading control) antibodies. The accumulation of the viral proteins 2C and 2BC reflects the level of replication.

Because the strength of the 2C-Arf1 interaction appears to correlate with BFA resistance, we compared the sensitivity of poliovirus and rhinovirus replication to BFA. HeLa cells were infected with an MOI of 10 of PV or rhinovirus 1A and incubated for 8 h in the presence of increasing concentrations of BFA. Analysis of the accumulation of viral proteins showed that poliovirus replication was blocked even at the lowest concentration of BFA (2μg/ml), while that of rhinovirus 1A was still robust even at the highest tested concentration (10 μg/ml) of the drug (Fig. 8D).

Thus, the increased interaction of 2C with Arf1 likely allows enteroviruses to tolerate the inhibition of GBF1 activity.

## Discussion

The antiviral therapeutics targeting host proteins hijacked by viruses are generally expected to pose a higher barrier to the development of resistance because the host proteins cannot change and because they support a process essential for the viral life cycle. Yet, the remarkable adaptability of enteroviruses repeatedly challenged this notion. To our knowledge, so far only an inhibitor (geldanomycin) of a cellular chaperone Hsp 90 required for the folding of enterovirus capsid proteins, and an inhibitor (E5) of an unidentified cellular factor supporting replication of diverse picornaviruses, proved to be refractory to the emergence of resistant mutants (60, 61). The low fidelity of enterovirus RNA-dependent RNA polymerase generates a cloud of related but not identical genomes (quasispecies) providing ample material for the selection of resistant mutants. Still, the probability of the immediate emergence of resistance rapidly diminishes if multiple mutations are required to overcome the selection pressure. The gradual selection of resistant genotypes with complex mutational landscapes is likely constrained by few evolutionary trajectories allowing additive accumulation of individual mutations (62, 63). Indeed, in this work, we document that resistance to high concentrations of BFA requires multiple mutations in the viral protein 2C, and the selection of such genotypes was only possible upon the gradual increase of the drug concentration, accompanied by step-by-step accumulation and disappearance of specific mutations. Thus, from the therapeutic standpoint, the development of viral resistance can be prevented or promoted by the particulars of the application protocol of an anti-viral drug.

While it is well documented that GBF1 is an essential cellular factor for enterovirus replication, the mechanistic understanding of why its activity is required for the assembly and/or functioning of the replication complexes is lacking. Our investigation of the replication of the mutant viruses highly resistant to BFA-mediated inhibition of GBF1 strongly suggested that the interaction of the viral protein 2C with activated Arf(s) may be required for replication. In cells infected with wt poliovirus, the supply of activated Arfs is normally ensured by the recruitment of GBF1 to the replication organelles through its interaction with the viral protein 3A (12, 34, 64). However, we observed that the replication of a BFA-resistant poliovirus mutant in the presence of BFA (when GBF1 is inhibited) was still accompanied by an accumulation of Arfs on the replication organelles. Since Arf association with membranes requires their activation, this raises two possibilities – either the mutant somehow restored the ArfGEF activity of GBF in the presence of BFA, or the Arfs associated with the replication organelles were activated by other, BFA-insensitive ArfGEF(s). Our data favor the latter possibility because the depletion of GBF1, or preventing its recruitment to the replication organelles (by incorporating a mutation in the viral protein 3A that severely restricts 3A-GBF1 interaction) had minimal effect on the BFA-resistant phenotype. The human genome encodes 12 proteins with Sec7 domains that are not inhibited by BFA. Some of them are expressed ubiquitously, while others have a more cell-type specific expression (reviewed in (30)). Our siRNA-mediated knockdown of individual ArfGEFs could not pinpoint a single BFA-insensitive ArfGEF that was essential for the replication of the BFA-resistant poliovirus mutant. Replication was most severely decreased by the siRNA-mediated depletion of Iqsec1/BRAG2, but significant decreases were also apparent by depleting Cytohesin1 and to a lesser extent some other BFA-insensitive ArfGEFs. Interestingly, the depletion of each of these BFA-insensitive ArfGEFs also inhibited the replication of wt replicon, suggesting that BFA-insensitive ArfGEFs (in addition to GBF1) play a currently unrecognized role in the replication process. These data are in line with our recent observation of the recruitment of a BFA-insensitive ArfGEF Cythohesin2/ARNO to the poliovirus replication organelles (65). 2C is one of the most conserved non-structural proteins of picornaviruses (66, 67). It is a multifunctional protein, a member of the helicase superfamily 3 (SF3) that has an ATP-dependent RNA helicase activity, and is important for RNA replication and virion assembly (68-72). Yet, the full spectrum of its functions in the viral replication cycle is far from understood. 2C is known to engage in interactions with multiple viral proteins (73-75), but only one cellular protein, reticulon3, a component of the ER membrane shaping complex has been reported as interacting with 2C (76). Herein, we describe a novel strong interaction between poliovirus 2C containing mutations conferring BFA-resistance with an active (GTP-bound) form of Arf1. Moreover, we detected an interaction between Arf1 and wild-type 2C of Rhinovirus 1A. Together, our data strongly suggest that the 2C-Arf interaction is conserved among enteroviruses, even though its strength may vary. The stronger rhinovirus 2C-Arf1 interaction may reflect the requirements of replication at significantly different temperature and/or cell type conditions than those for enteroviruses– upper respiratory tract and alimentary tract, respectively. Interestingly, it was reported that the 3A proteins of rhinovirus 2 and 14 do not strongly interact with GBF1. This was attributed to the differences in the N-terminal sequences between poliovirus and rhinovirus 3As (77). Thus, the stronger 2C-Arf1 interaction could provide rhinoviruses with an alternative, GBF1-independent, recruitment of Arf1 to the replication organelles. Consistent with the model that BFA resistance is based on increased 2C-Arf1 interaction, replication of rhinovirus 1A was much less sensitive to BFA inhibition than that of poliovirus.

The mutations increasing interaction of poliovirus 2C with Arf1 were concentrated in the N-terminal part of the protein which is not resolved in the crystal structure, but is known to contain membrane targeting and RNA binding motifs (78-81). It was proposed that ATPase, RNA-binding, and RNA helicase activities of 2C require the formation of different homo-oligomers, from 2 to 8 subunits, with the hexamer modeled on the SV40 large T antigen, another member of SF3 family helicases, thought to be the likely configuration of the protein in the replication complexes (80-83). In this model, the N-terminal part of the protein would be facing the membrane surface where it will be accessible to interacting with active membrane-bound Arfs. The essential role of host factors in 2C activity is supported by a significant variation of replication of the same BFA-resistant mutant, i.e. containing the same mutations in 2C, in different cell lines.

The data reported in this paper and our previous observations (33) indicate that in infected cells at least two different subpopulations of 2C exist – those that are located in Arf-enriched and Arf-depleted domains, and that active Arfs may stabilize and/or activate the conformation of 2C required to support RNA replication. This also implies that the functionalities of Arf-enriched and Arf-depleted 2C subpopulations would be different. Thus, the interactions with cellular components in spatially distinct domains may increase the spectrum of activities performed by viral proteins, effectively compensating for the limited coding capacity of the enterovirus genome.

Collectively, our results demonstrate that the inhibitory effect of BFA on enterovirus replication can be reversed by mutations that increase the interaction of the viral protein 2C with the active (GTP-bound) form of Arf1, which strongly suggests that the requirement for such interaction determines the sensitivity of enterovirus infection to BFA in the first place. Thus, at least a subpopulation of 2C on the replication membranes likely behaves like an Arf effector protein, with Arf1 binding triggering/stimulating/initiating specific function(s) of 2C in the replication process. Our data also document that the development of viral resistance to the inhibition of a cellular protein exploits pre-existing pathways and interactions with cellular factors to restore the essential functions inaccessible to the wt virus in the presence of the inhibitor, rather than the establishment of a new mode of replication. In this regard, it appears that the 2C-Arf(GTP) interaction rather than GBF1 activity is indispensable for enterovirus replication, which may guide future development of effective anti-virals.

## Figure legends

**Figure S1**. A summary of the mutations observed in the 2C-3A region of individual viral isolates and after sequencing PCR fragments obtained from the total viral RNA at sequential selection steps (10 passages at 4 μg/ml of BFA followed by 10 passages at 20 μg/ml of BFA followed by 25 passages at 50 μg/ml of BFA). Green highlights indicate resistant genotypes taken for further analysis.

**Figure S2**. Replication of Renilla polio replicons with the indicated combinations of individual mutations in 2C in the presence of 25 μg/ml BFA. Relative replication is calculated for the signal up to 8 h post-transfection (dashed line) and is normalized to the replication of the replicon with the full complement of resistant mutations in 2C (59/2C).

**Figure S3. A**. Replication of the polio Renlla replicon with BFA-resistant mutations in 2C (59/2C) in cells treated with siRNAs targeting individual BFA-insensitive ArfGEFs for 72 h in the presence of 25 μg/ml of BFA. Relative replication is calculated for the signal up to 8 h post-transfection (dashed line) and is normalized to the replication of in cells treated with control siRNA. **B-E**.

