## Supplementary figures and images for "The Development of Resistance to an Inhibitor of a Cellular Protein Reveals a critical interaction between the enterovirus protein 2C and a small GTPase Arf1"

### Supplementary Figure 1

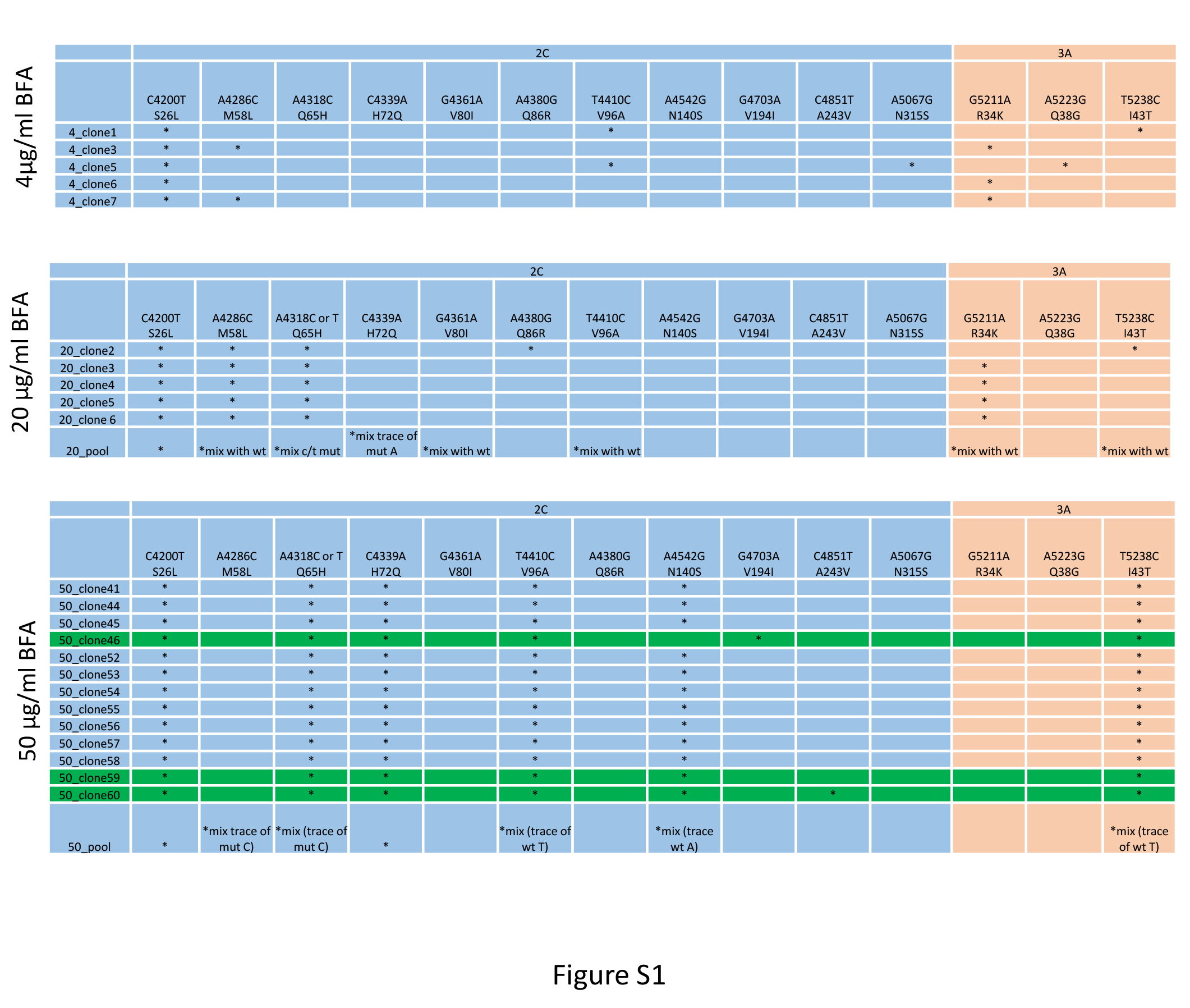

### Supplementary Figure 2

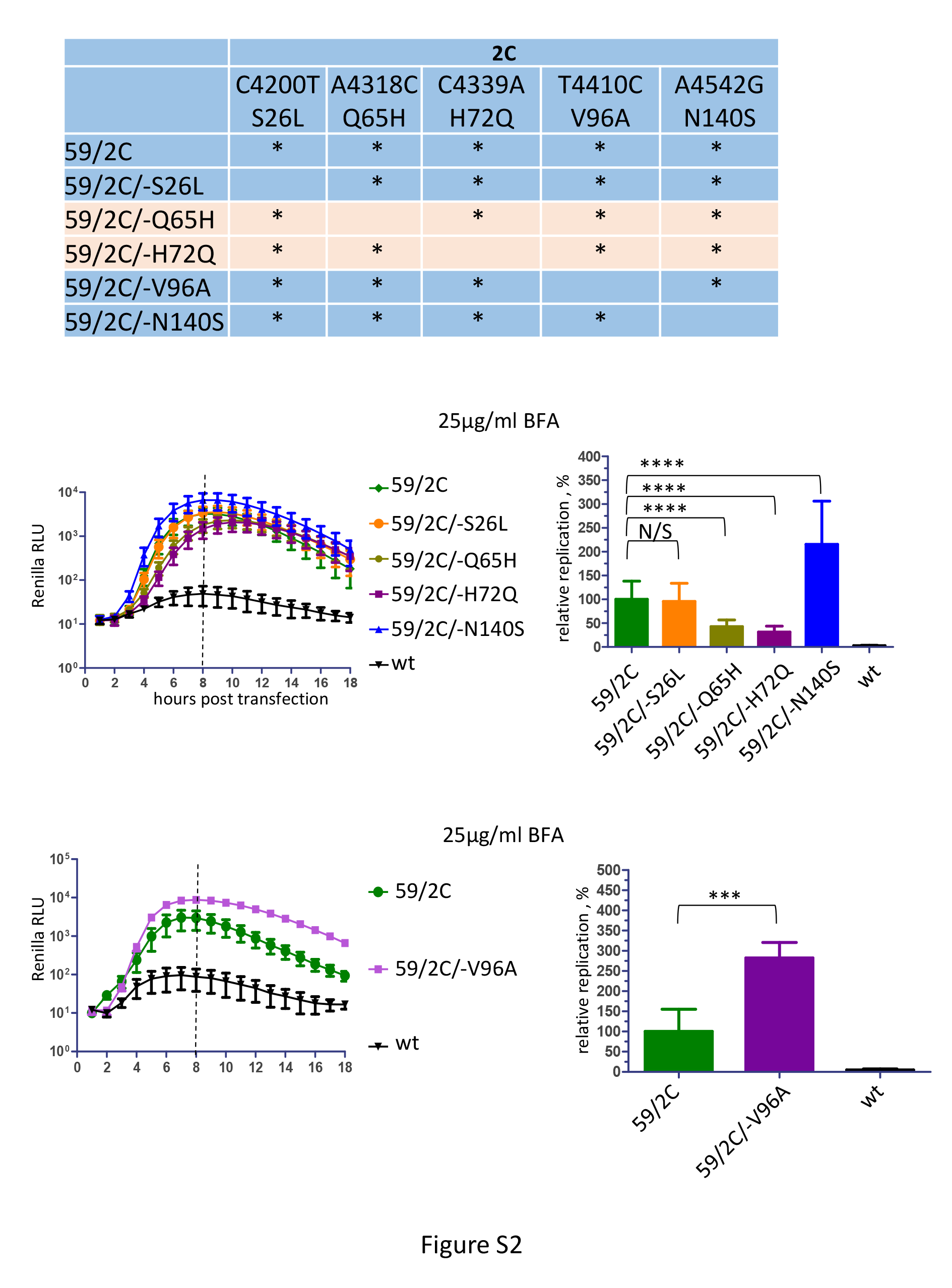

### Supplementary Figure 3

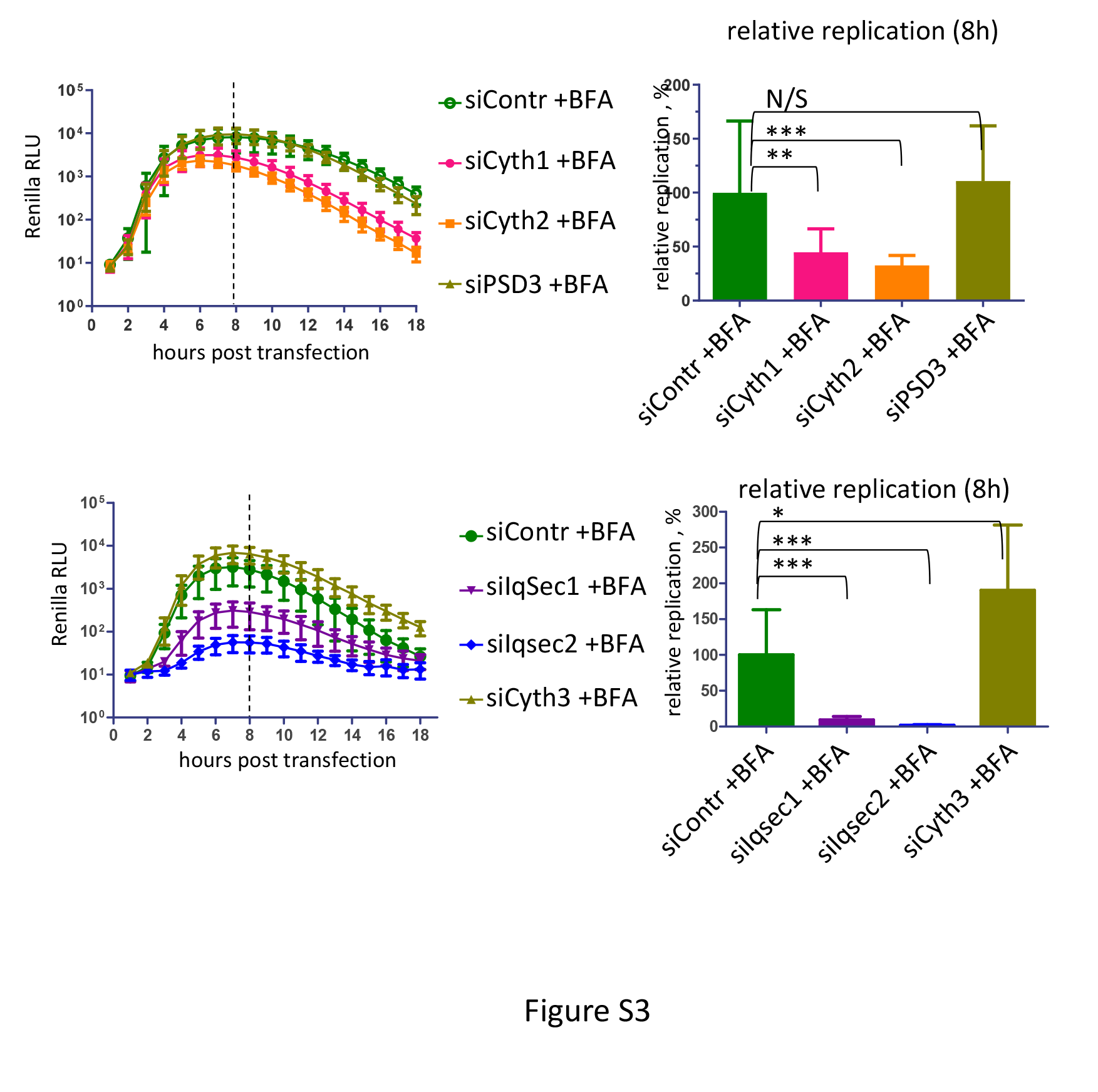
